# Host-microbiota interactions contributing to the heterogeneous tumor microenvironment in colorectal cancer

**DOI:** 10.1101/2023.07.17.549261

**Authors:** Xiaoyi Li, Dingfeng Wu, Qiuyu Li, Jinglan Gu, Wenxing Gao, Xinyue Zhu, Wenjing Yin, Ruixin Zhu, Lixin Zhu, Na Jiao

**Affiliations:** Department of Nephrology, Children's Hospital, Zhejiang University School of Medicine, National Clinical Research Center for Child Health, Hangzhou 310058, Zhejiang, P.R. China.; The Shanghai Tenth People's Hospital, School of Life Sciences and Technology, Tongji University, Shanghai 200072, P. R. China.; Department of Colorectal Surgery, Guangdong Institute of Gastroenterology, Guangdong Provincial Key Laboratory of Colorectal and Pelvic Floor Diseases, The Sixth Affiliated Hospital, Sun Yat-sen University, Guangzhou 510655, P.R. China.

**Keywords:** Colorectal cancer, Transcriptomic, Microbiome, Host-microbiota interactions, Ferroptosis

## Abstract

**Background:** Colorectal cancer (CRC) is a highly heterogeneous cancer with four consensus molecular subtypes (CMSs) characterized by distinct tumor microenvironment (TME). We aimed to depict the characteristics of host-microbiota interactions and their contributions to TME in each CMS.

**Methods:** Host transcriptome and intratumoral microbiome profiles of 594 CRC samples were derived from RNA-Seq data from TCGA. Differential host genes and microbes among CMSs were identified. Immune microenvironments were assessed by CIBERSORTx and ESTIMATE and microbial co-abundance analyses were performed by FastSpar. Host-microbiota associations were evaluated by LASSO penalized regression in each CMS.

**Results:** Along with distinct host gene signatures, including ferroptosis-related genes and immune microenvironments, 293, 153, 66 and 109 intratumoral differential microbial genera were identified within each of the four CMSs, respectively. Furthermore, the host-microbiota interactions contributed to distinct TME in each CMS, represented by 829, 1,270, 634 and 1,882 robust gene-microbe associations, respectively. The TME in CMS1 was featured with inflammation-related HSF1 activation and interactions between genes of endothelin pathway and *Flammeovirga*. Integrins-related genes positively correlated with *Sutterella* in CMS2 while CMS3 displayed microbial associations with biosynthetic and metabolic pathways. Genes in collagens biosynthesis positively correlated with *Sutterella*, contributing to homeostasis disturbance in CMS4. Besides, ferroptosis dysregulation was more remarkable in immune-high subtypes, which might partly result from the colonization of tissue microbes.

**Conclusions:** We systematically profiled the landscapes of TME, of each CMS in CRC, encompassing host genes, intratumoral microbiome and their interactions, which could illuminate novel mechanisms for the heterogeneity in CRC and potential therapeutic targets.

## Introduction

Colorectal cancer(CRC) is one of the most common cancers worldwide with high incidence and mortality, reaching 1.9 million new cases and 935,000 deaths annually(1). CRC is highly heterogeneous at the molecular level due to genetic, epigenetic, and tumor microenvironment(TME) alterations, which result in variable clinical outcomes and treatment responses(2, 3). TME not only consists of tumor-infiltrating cells such as tumor-associated macrophages(TAMs), B cells, and T cells, but the whole surrounding environment of the tumor, including vasculatures, extracellular matrixes(ECMs), and other constituents, such as microbiota(4). To better stratify CRC patients, the international consortium of colorectal cancer subtyping(CRCSC) defined a system based on tumor transcriptome and characterized four consensus molecular subtypes(CMS1-4) with molecular and clinical differences: CMS1(MSI), CMS2(canonical), CMS3(metabolic) and CMS4(mesenchymal)(5). Besides, different clinicopathologic properties were observed among CMSs, for example, the HSP90 inhibition could effectively alleviate the chemoresistance for CMS4 patients(6). Gene expression profiles of CMS2 were largely influenced by DNA copy number gains in malignant cells, whereas the profiles of CMS1 and CMS4 were driven by the infiltrated non-malignant cells in the TME(7). Moreover, the state-of-the-art approach, single-cell transcriptomics elucidated the cellular diversity within the TME across CMSs, highlighting the diverse phenotypes of cancer-associated fibroblasts(CAFs) and TAMs(8).

In recent years, emerging evidence has highlighted the vital roles of the microbiome in detecting CRC and mediating the potential carcinogenesis of CRC, which was probably through the production of microbial metabolites that interacted with the host-immune system(9–14). Pioneering researches, including ours, have identified adenoma-specific bacterial biomarkers for early detection of CRC and bacteria-fungi interactions promoting the development of CRC based on fecal microbiome sequencing(15–19). The remarkable work by Poore et al. demonstrated the high rates of microbial sequencing reads in tumor tissues derived from host RNA-Seq data and identified unique microbial signatures within and between most major types of cancer(20), which paved the way for investigating the roles of intratumoral microbes in cancer pathogenesis. Since microbiota is now regarded as a crucial component of TME, recent efforts have revealed CMS associated microbes and potential mechanisms. For example, CMS4 patients with a high abundance of *Fusobacterium nucleatum* or *Fusobacteriales* in tumor tissues were associated with worse clinical outcomes and severe inflammatory responses(21) while CMS1 patients were related to *Parvimonasmicra*(22). Besides, both Purcell et al and Visnovska et al described the microbial patterns in different CMSs based on 16S rRNA sequencing(23, 24). However, limited studies explored the whole landscape of gut microbiota and their interplays with host genes and how they contribute to the heterogeneity of CMSs remain unresolved.

Therefore, this study aimed to characterize the global features of TME focusing on host-microbiota interactions in CMSs and their potential implications in CRC pathogenesis. In this study, we collected the host gene expression profile and corresponding gut microbial abundance matrix of the TCGA-CRC samples, identifying CMS-specific host genetic and microbial signatures. Further, CMS-specific and shared host-microbiota interactions were profiled in each CMS, contributing to the understanding of the TME heterogeneity of CMSs and potential strategies for personalized intervention.

## Methods

### 1. Resource Availability

#### 1.1 Materials availability

This study did not generate new unique reagents.

#### 1.2 Data and code availability

All the data for this manuscript are publicly available. The pre-processed host mRNA expression profile data in this study have been deposited in the TCGA database. The pre-processed microbial abundance profile data and corresponding metadata are available at http://ftp.microbio.me/pub/cancer_microbiome_analysis/. The survival phenotypic data of TCGA-COAD and TCGA-READ samples are accessible through the SAGE Synapse platform under accession syn7343873. All programming scripts can be found at our GitHub repository link: https://github.com/FragmentsLi/CMS.

### 2. Study design and data preprocessing

Since this study focused on the heterogeneity of different CRC subtypes, we began with 618 CRC samples with matched microbial abundance and host gene expression profiles. After excluding samples without survival information or with an overall survival(OS) time of 0, 594 samples remained.

In addition, we removed 12 genera with raw microbial abundance equal to 0 in all CRC samples from the normalized microbial profiles to avoid false positives. ComBat was applied to remove batch effects of sequencing platforms in host gene expression data(25). Principal component analysis(PCA) was performed to visualize the sample distribution before and after batch correction(See Supplementary Figure 1).

### 3. Host gene expression characterization analysis

#### 3.1 CMS classification

The CMS classification labels of CRC samples were obtained from the SAGE Synapse platform(Accession: syn2623706)(5). Samples without labels were classified using the random forest(RF) classifier in R package “CMSclassifier”(v 1.0.0)(5) based on the preprocessed host gene expression data.

#### 3.2 Immune cell infiltration analysis

CIBERSORT(v 1.03) was performed to estimate the infiltration of 22 immune cell types among CMSs based on the preprocessed host gene expression data(26). Similarly, the Estimation of Stromal and Immune cells in Malignant Tumours using Expression data(ESTIMATE) algorithm(v 1.0.13) was used to evaluate the tumor purity scores, stromal cells, and immune cells infiltration levels among CMSs(27). The infiltration results across CMSs were considered statistically significant when *P* value<0.05, by the Kruskal–Wallis rank sum test.

#### 3.3 Differential expression analysis

To identify robust differential genes, we first selected effective genes from the entire list of the preprocessed host genes by (1)excluding non-protein-coding genes via R package “biomaRt”(v 2.46.3)(28); (2)excluding genes expressed in less than half of all the CRC samples; and (3)excluding genes with variances < 25% quantile of variances across all samples. We obtained a total of 11,304 effective genes, which were further analyzed to determine differential genes among CMSs by implementing Kruskal-Wallis rank sum test followed by Dunn’s *post hoc* test for multiple comparisons. Differential genes in one specific CMS were defined with Kruskal-Wallis test *P* value <0.05 and Dunn’s *post hoc* test *P* value<0.05 in all comparisons. Pathway enrichment analysis was performed based on the differential genes in each CMS via clusterProfiler(v 4.2.2) with a threshold *P* value <0.05.

#### 3.4 Characterization of ferroptosis-related genes

Ferroptosis-related gene lists were obtained from FerrDB V2(http://www.zhounan.org/ferrdb/)(29). This list of 564 genes includes 9 marker genes, 264 driver genes, 238 suppressor genes, and 110 unclassified genes. These genes were further mapped to differential genes to investigate the pathological roles of ferroptosis in CRC.

### 4. Gut microbiome analysis

#### 4.1 Microbial ecological analysis

To measure the diversity and richness of microbial communities, the preprocessed microbial abundance profiles were transformed to non-negative data by nneg function in R package “NMF”(v 0.21.0)(30). Alpha diversity metrics were estimated via Shannon and Gini Simpson indices for each sample, and similarly, Kruskal–Wallis rank sum test was used to assess the differences among CMSs(*P* value<0.05). The microbial community difference was measured by beta diversities based on Bray-Curtis distances, and the differences among CMSs were tested using Analysis of similarities(ANOSIM) by 9,999 permutations in vegan packages(v 2.6.2).

#### 4.2 Differential abundance analysis

To identify differential microbes, differential abundance analysis was performed among CMSs by implementing Wilcoxon rank sum test against 797 genera with prevalence over 20% in all samples. Differentially abundant genera were defined as they showed significant statistical differences between a specific CMS and all other CMSs combined (*P* value<0.05).

#### 4.3 Microbiotaco-abundance analysis

Microbial co-abundance associations were inferred based on the microbial counts in each CMS through FastSpar(v 0.0.10)(31), a rapid C++ implementation of SparCC algorithm. In FastSpar, 1,000 iterations, 100 exclusion iterations, and 1,000 random permutations were used to calculate *P* values. The correlations were considered statistically significant with *P* value<0.05 and absolute correlation *r*>0.7, which were further visualized through Gephi(v 0.9.7).

### 5. Host-microbiota association analysis and pathway enrichment

#### 5.1 Procrustes analysis and mantel test

Procrustes analysis and mantel test were performed toassess the structural similarity between host gene expression and microbial abundance profiles. The host gene expression profiles were measured with Aitchison’s distance and microbial abundance profiles with Bray-Curtis distance through R package “vegan”(v 2.6.2). The statistical significance of the correspondence was measured by 9,999 permutations.

#### 5.2 Host-microbiota association analysis

To identify specific robust associations between host genes and gut microbes, a LASSO penalized regression model developed by Priya et al(32) was implemented to identify host-microbiota associations, i.e., gene-microbe associations, in each CMS. First, in the model estimation, the expressions of the differential genes of each sample in a CMS served as responses while their paired microbial abundances of the differential microbes served as predictors. The desparsified LASSO achieved 95% confidence intervals and *P* values for the coefficient of each predictor (microbe) corresponding to the given response(host gene). The Benjamini-Hochberg(FDR) method was used to adjust for multiple tests. Next, stability selection based on a 100 times re-sampling was performed for each host gene to select considered stable correlated microbes which appeared in at least 60% of the fitted models. Finally, the overlapped between gene-microbe associations with FDR<0.1 and gene-microbe associations considered stable were identified as significant robust gene-microbe associations. In addition, ferroptosis related gene-microbe associations were selected from significant robust gene-microbe associations with the criterion of ferroptosis related differential genes having an absolute of Spearman’s rho coefficient> 0.3 calculating between gene expressions and paired microbial abundances. All gene-microbe association networks were visualized through Cytoscape(v 3.9.1).

#### 5.3 Pathway enrichment

Genes in significant robust gene-microbe associations in each CMS were subjected to enrichment analysis based on Kyoto Encyclopedia of Genes and Genomes(KEGG), Pathway Interaction Database(PID) and REACTOME gene sets from the MsigDB(33). Significant host pathways were determined by Fisher’s exact test with *P* values<0.05, where the input of the LASSO models(i.e., differential genes in one specific CMS) served as background genes and genes in significant robust gene-microbe associations served as genes of interest.

## Results

### 1. CMS-specific characteristics of host gene expression

To comprehensively explore the TME characteristics of different CMSs in CRC, we performed analyses on host gene expression patterns, intratumoral microbial signatures, and host-microbe associations(Figure 1). After quality control, a total of 594 CRC samples with matched host gene expression profiles and intratumoral microbial abundance profiles were extracted from the TCGA database and then classified into four CMSs based on host gene expression data, excluding sample with mixed signatures(Figure 1). Of all metadata variables, we found that CMS1 samples were predominantly located on the right side of the abdomen(count, percentile: 49, 64.5%) while CMS2 were on the left side of the abdomen(count, percentile:139, 62.6%, See Supplementary Table 1) as reported in previous studies(34–36). The different distribution might be attributed to the different incidences of MSI-status, *BRAF* mutation and the *TP53* mutation rates(37). In addition, significant associations between CMS classification and either TNM or tumor stages were identified(chi-square test, *P* value<0.05). Samples in CMS2 and CMS4 have a higher possibility of metastasis with cancers in lymph nodes and distant metastasis than other CMSs(See Supplementary Table 1).

**Figure 1.**
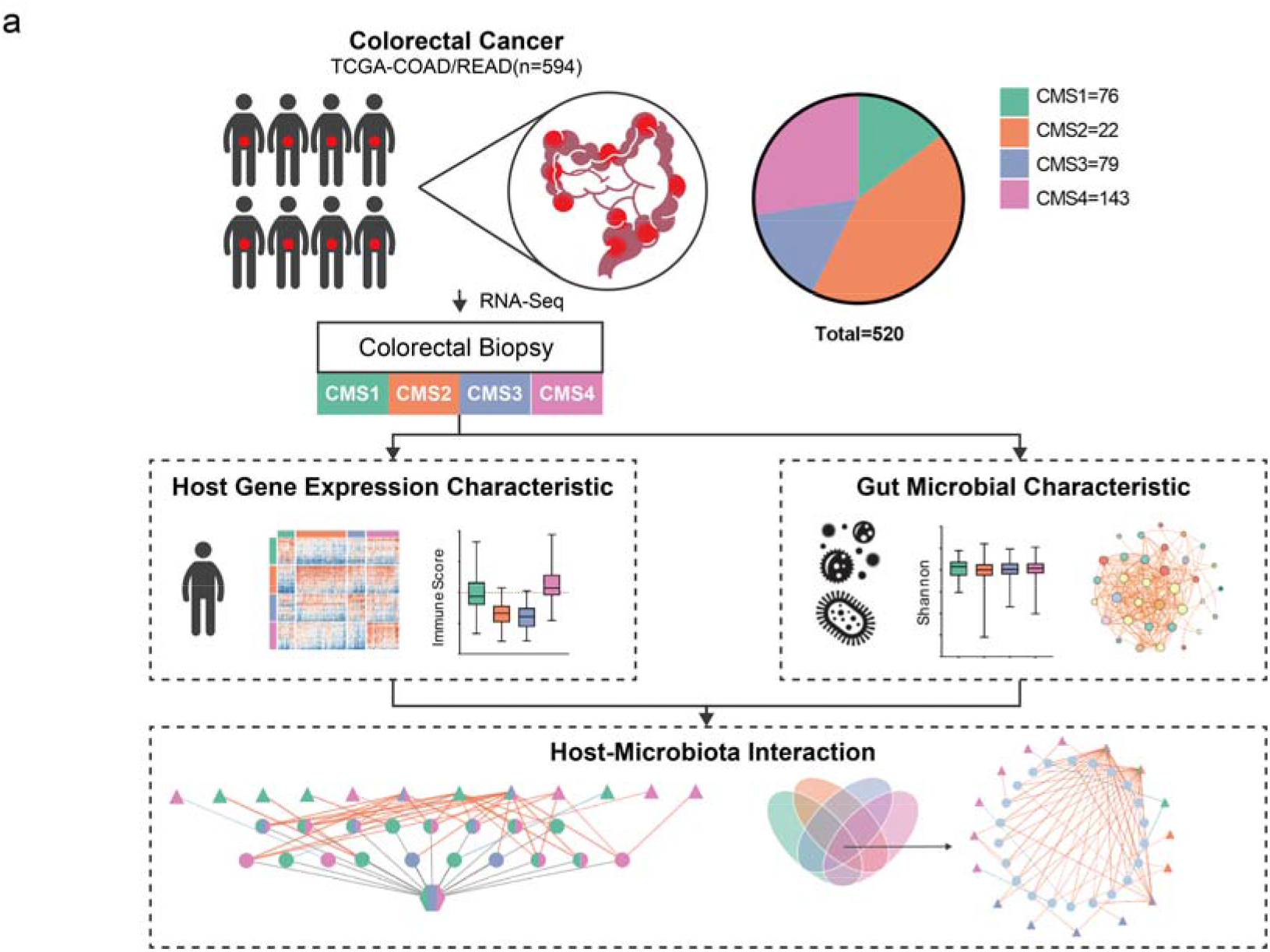
Workflow of the study. Paired host gene expression data and gut microbial abundance data of 594 CRC samples from TCGA were classified into four CMSs based on gene expression profiles. Characteristics of the host gene expression and gut microbiota were examined in each CMS, followed by host-microbiota interaction analysis with the integrated data.

Though the molecular heterogeneity of four CMSs against host transcriptome has been comprehensively clarified(5, 38–40), here we revisited the host gene expression profiles and consistently found a large number of differential genes (2,403 in CMS1(See Supplementary Table 2a), 1,140 in CMS2(See Supplementary Table 2b), 1,359 in CMS3(See Supplementary Table 2c) and 2,091 in CMS4(See Supplementary Table 2d)), highlighting the great heterogeneities among four CMSs(Figure 2a). Since the vital roles of ferroptosis have been reported on the progression and treatment of CRC(41–44), we examined the expression patterns of ferroptosis-related genes in four CMSs and identified 89, 34, 41, and 76 ferroptosis-related differential genes in CMS1, CMS2, CMS3, and CMS4, respectively(Figure 2b). Moreover, the expression levels of these ferroptosis-related genes highly varied among CMSs with no common driver and suppressor gene identified(Figure 2b). Specifically, in CMS1, a series of ferroptosis-related differential genes exhibited strong positive associations internally(See Supplementary Figure 2), including up-regulated driver genes *IDO1*, *TRIM21*, *SLC25A28* and down-regulated suppressor genes *PARP9*, *PARP12*, *PARP14*, and *SREBF1*. Fewer ferroptosis-related differential genes were identified in CMS2, with driver gene *BRD7* and suppressor gene *ETV4* up-regulated, while another suppressor gene *ARF6* down-regulated(See Supplementary Figure 3). Interestingly, CMS3 was probably negative for ferroptosis, which was reflected by the down-regulation of a panel of positively associated genes, including driver genes *WWTR1*, *DDR2*, and *ZEB1*(See Supplementary Figure 4). Most of the ferroptosis-related genes were positively associated with CMS4(See Supplementary Figure 5). These genes included up-regulated driver genes *TGFBR1*, *WWTR1*, *DDR2* and down-regulated suppressor genes *EZH2*, *FANCD2*, and *CDC25A*. Taken together, these findings suggest differential activities and patterns of ferroptosis among the CMSs, and that the ferroptosis status was suppressed in CMS3 but relatively active in CMS1 and CMS4.

**Figure 2.**
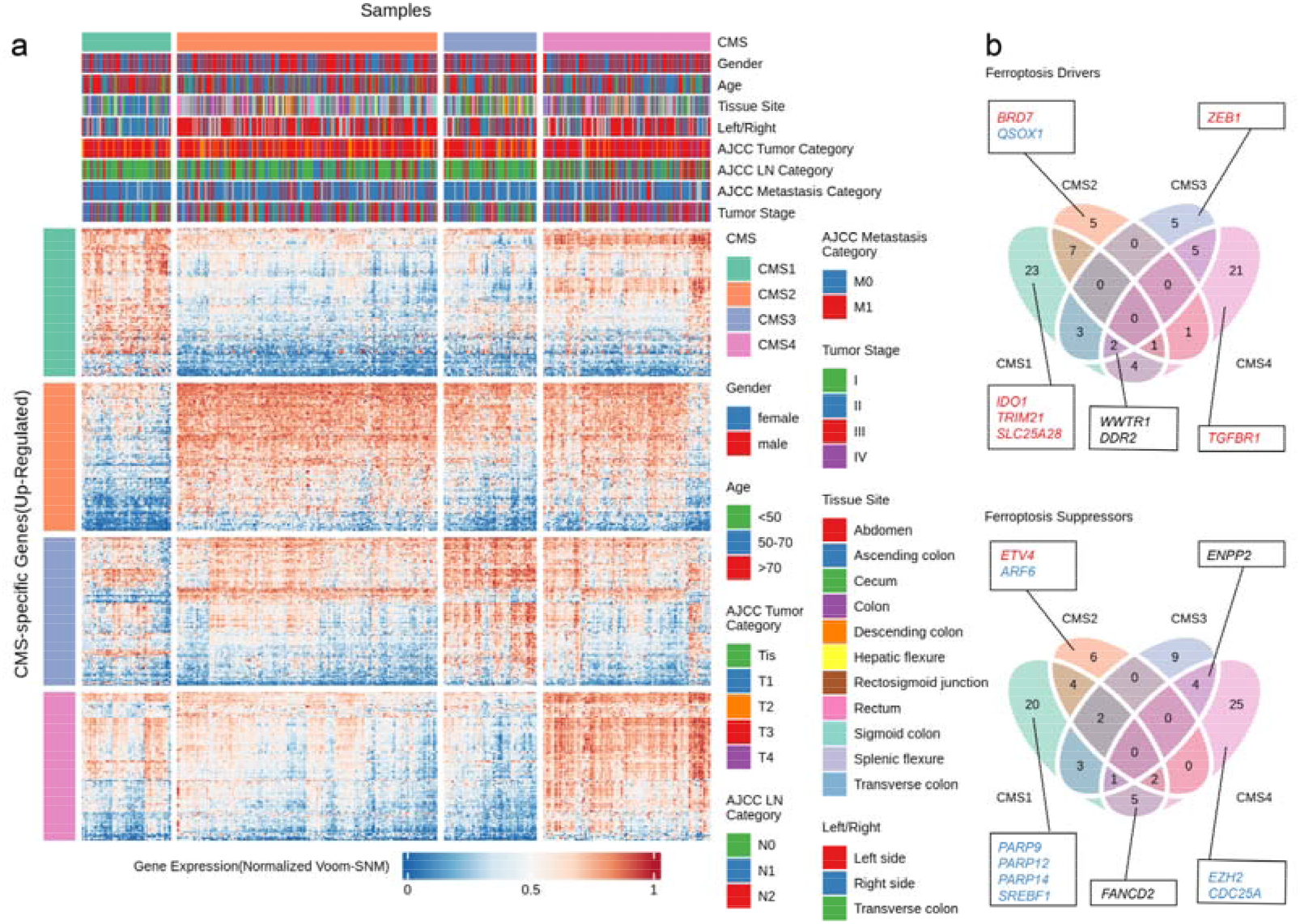
CMS-specific characteristics of host gene expression. **(a)**Heatmap displaying the top 100 up-regulated differential genes among CMSs. Samples and genes were clustered within CMS, indicated by the colors. The clinical characteristics of samples were indicated on the top. Genes ranked in the top 100 according to the maximum differences between any specific CMS and the others were selected. **(b)**Venn diagram of the ferroptosis related drivers and suppressors across CMSs, significant ferroptosis related genes were annotated. For genes in one specific CMS, up-regulated genes were colored in red while down-regulated genes were colored in blue. Genes that appeared in multiple CMSs were colored in black.

We then performed pathway enrichment analysis on differential genes of each CMS to elucidate their biological functions. In total, we identified 77 pathways in CMS1, i.e., CMS1-specific pathways(See Supplementary Table 2e), 19 CMS2-specific pathways(See Supplementary Table 2f), 44 CMS3-specific pathways(See Supplementary Table 2g) and 66 CMS4-specific pathways(See Supplementary Table 2h). Among them, most of the CMS1 and CMS4-specific pathways were related to immune activation. Furthermore, we explored the heterogeneity of TME by estimating the immune-stromal score and the proportion of immune cells in TME for each CMS. Firstly, CMS1 and CMS4 showed higher immune scores than CMS2 and CMS3(Figure 3a). Similarly, CMS4 displayed the highest stromal score, suggesting the presence of a large number of stromal cells including mesenchymal stromal cells (MSCs). Then, with CIBERSORTx, the proportions of 22 immune cells were estimated and manifesting distinct immune microenvironments across CMSs. In detail, CMS1 exhibited higher levels of CD8+ T cells, follicular helper T cells, and M1 macrophages as well as decreased Treg cells and monocytes compared to the others (Figure 3b). In contrast, CMS4 was dominated by M0 and M2 macrophages accounting for over 20% of total cells, with fewer M1 macrophages and CD8+ T cells(Figure 3b), reflecting a distinct immune pattern probably due to higher infiltration of inflammatory components in CMS4(45). Meanwhile, CMS2 and CMS3 exhibited higher fractions of plasma cells compared to CMS1 and CMS4 (Figure 3b).

**Figure 3.**
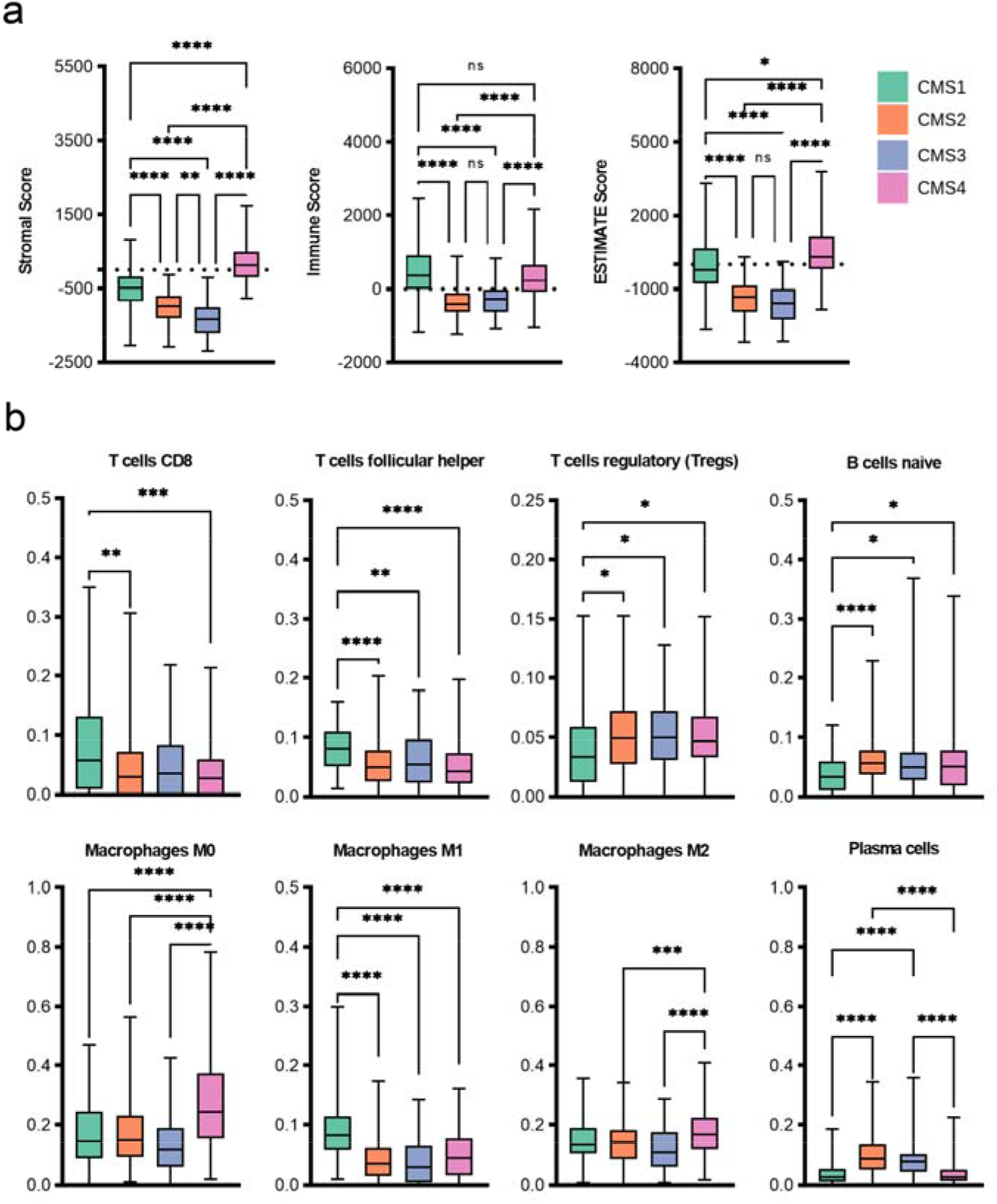
Immune characteristics of CMSs. **(a)**Differences in immune scores, stromal scores, and tumor purity calculated by ESTIMATE among CMSs. **(b)**Differences in immune cell infiltration were calculated by CIBERSORTx among CMSs. See the differences in the infiltration of the remaining 14 immune cells in Supplementary Figure 6.

### 2. CMS-specific characteristics of gut tissue microbiota

Intratumoral microbiota emerges as an important factor that contributes to the heterogeneity of TME(13), we thus examined the characteristics of tumor-associated microbiota in different CMSs at multiple layers based on the microbial profiles derived from Poore’s work(20), which has been well annotated after decontamination and benchmarking. First, though decreased microbial alpha diversity was reported in CRC tumor tissues compared toadjacent normal tissues(46), we didn’t observe asignificant difference in microbial diversity among CMSs and the intratumoral microbiota generally displayed similar compositions (Figure 4a, See Supplementary Figure 7), reflecting a considerable difference between intratumoral and fecal microbiota, which was further validated in the microbial composition at the phylum level. Proteobacteria was the dominant phylum accounting for 38.7-38.8% relative abundance in CRC tissues, while the dominant fecal phyla Firmicutes and Bacteroidetes only represented 13.8-13.9% and 9.2%, respectively, of the fecal microbiota(Figure 4b). In addition, Euryarchaeota (3.9%), a major phylum from archaea, and virus(8.8%) were also detected in tumor tissues.

**Figure 4.**
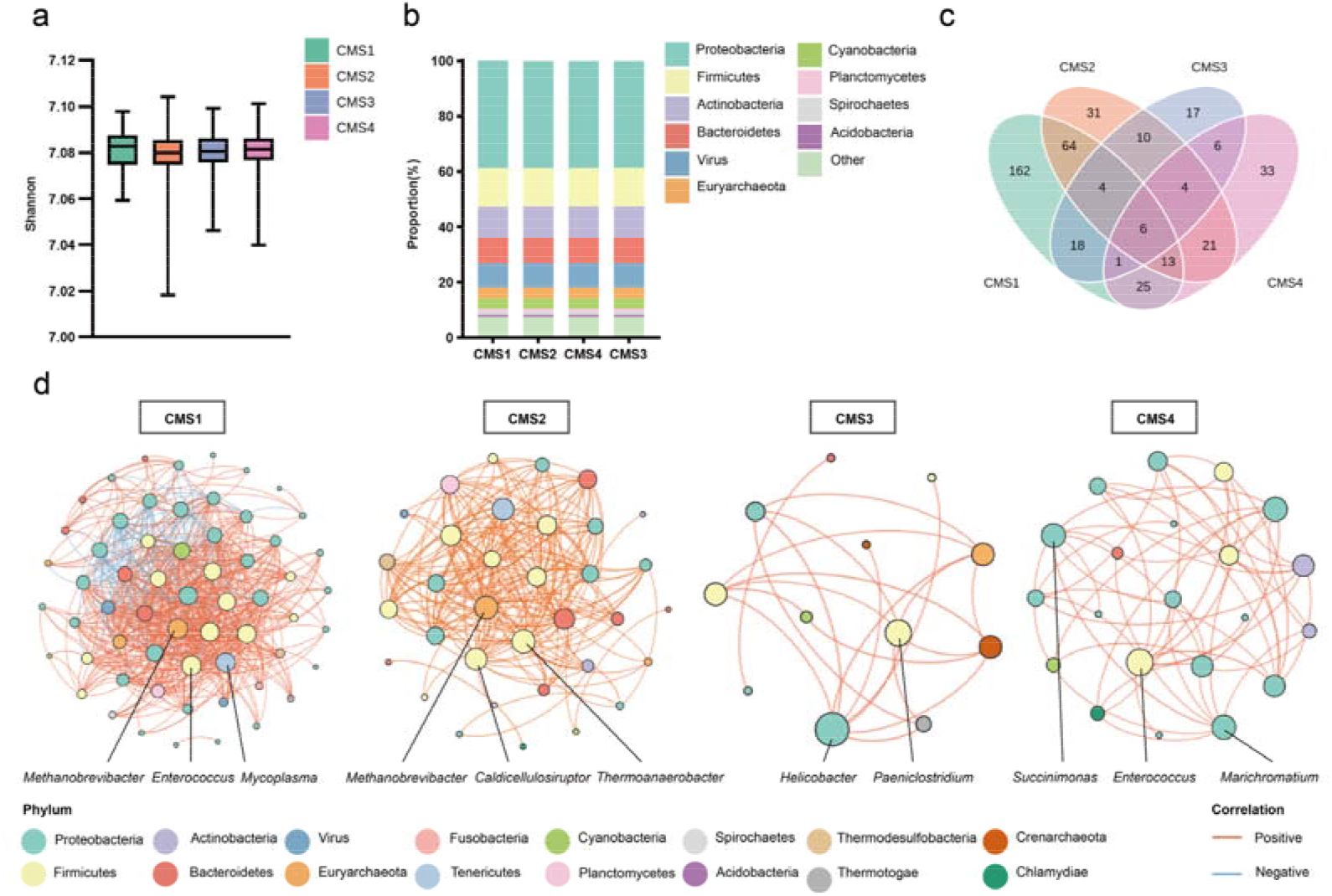
CMS-specific characteristics of intratumoral microbiota. **(a)**Alpha diversities(Shannon index) in the CMSs. See the Gini Simpson indices and beta diversities in the CMSs in Supplementary Figure 7. **(b)**The top 10 abundant phyla varied among CMSs. **(c)**Venn diagram of the CMS-specific genera counts among CMSs. **(d)**CMS-specific co-abundance networks. Only significant correlations(absolute correlations *rho*> 0.7) are shown, each node indicates one genus. The colors of nodes represent the phyla to which the genus belongs. The colors of the edges represent the positive(orange) or negative(blue) correlation between genera.

At the genus level, we identified considerable differential genera among CMSs, including 293 differential genera in CMS1, 153 in CMS2, 66 in CMS3, and 109 in CMS4(Figure 4c). There were 195 genera that were significantly increased in CMS1(See Supplementary Table 3a), accounting for 66.6% total relative abundance, and most of them belong to Proteobacteria including *Desulfuromonas, Bilophila*, and *Nitrosomonas*. Besides, some SCFA-producing genera also increased in CMS1, such as *Akkermansia*, *Lachnoclostridium*, and *Ruminococcus*. On the other hand, 98 genera exhibited decreased abundances in CMS1, including *Paeniclostridium* and *Marinitoga*. In CMS2, increased abundances were observed for 98 genera(See Supplementary Table 3b), including *Paenarthrobacter*, *Lactobacillus*, and *Thermoanaerobacter,* while decreased abundances were observed for 55 genera, including SCFA-producers *Dorea*, *Ruminococcus*, and *Butyrivibrio*. In CMS3, 33 genera were enriched, such as *Ruminococcus*, *Rhodothermus*, and *Bacteroides*, while the abundances of *Sutterella*, *Collimonas* and *Lactobacillus* were reduced(See Supplementary Table 3c). In CMS4, 61 genera were enriched, including *Sutterella*, *Actibacterium*, *Enterococcus*, and *Flammeovirga*. Meanwhile, 48 genera were decreased, including *Desulfococcus*, *Fusobacterium*, and *Succinimonas*(See Supplementary Table 3d). These findings suggest that immune-high subtypes(CMS1 and CMS4) exhibited an opposing abundance pattern in intratumoral microbiota against the immunosuppressive subtypes(CMS2 and CMS3), with top enriched genera in immune-high subtypes such as *Flammeovirga*, *Sutterella*, and *Collimonas* depleted in immunosuppressive subtypes. Besides, we also identified differential bacteriophages in the CMSs, which may have contributed to the changes in the bacteriome.

Next, co-abundance associations of the differential genera were examined in each CMSs(Figure 4d). Distinct patterns were observed for different CMSs. Compared to the others, CMS1 exhibited more complex associations with 605 edges among 60 genera, consistent with its strong immune activation status in which gut microbiota could take part(See Supplementary Table 3e).*Methanobrevibacter*, *Enterococcus*, and *Mycoplasma*, with depleted abundance, were highly connected to other nodes in the network(higher degrees), suggesting their hub roles in maintaining the microbial microecology in CMS1 network. The co-abundance network of CMS2 consisted of 34 genera with 211 edges, including increased *Methanobrevibacter*, *Thermoanaerobacter*, and *Caldicellulosiruptor*(See Supplementary Table 3f). The co-abundance network in CMS3 was the sparsest(21 co-abundance associations among 12 genera, see Supplementary Table 3g), with highly connected nodes *Helicobacter* and *Paeniclostridium*. In CMS4, there were 53 co-abundance correlations among 21 genera, with *Enterococcus*, *Marichromatium*, and *Succinimonas* as the hub genera in the co-abundance network (See Supplementary Table 3h). *Succinimonas* was less abundant in CMS4 and was previously reported as a biomarker in recurrence or metastasis of lung cancer(47). Taken together, our data displayed distinct structures and key genera of intratumoral microbial environments among CMSs, suggesting potential contributions of intratumoral microbes in the heterogeneity of TME.

### 3. Host-microbiota interactions in CMSs

To understand the host-microbiota interactions in different CMSs, we performed an integrated analysis to identify associations between host gene expression levels and microbial genera. By Procrustes analysis and mantel test(See Supplementary Figure 8), 829, 1,270, 634, and 1,882 robust host gene-microbe associations were identified in CMS1, CMS2, CMS3, and CMS4, respectively. These represent associations between 717 host genes and 180 microbial genera in CMS1(See Supplementary Table 4a), between 814 host genes and 100 microbial genera in CMS2(See Supplementary Table 4b), between 547 host genes and 59 microbial genera in CMS3(See Supplementary Table 4c), and between 1,313 host genes and 89 microbial genera in CMS4(See Supplementary Table 4d).

To elucidate the biological functions mediated by host genes that are associated with intratumoral microbiota across CMSs, we subjected the host genes from gene-microbe associations in each CMS to perform pathway enrichment analysis(Figure 5a). Overall, we identified 21 pathways enriched by host genes from gene-microbe associations in CMS1(See Supplementary Table 4e), 6 pathways in CMS2(See Supplementary Table 4f), 18 pathways in CMS3(See Supplementary Table 4g), and 29 pathways in CMS4(See Supplementary Table 4h), including several CMS-specific pathways(Figure 5a). For a better understanding of the potential mechanism mediated by host gene-microbiota associations, we constructed the interaction network among host genes, gut microbiota, and the enriched pathways (Figure 5b). Host genes from gene-microbe associations in CMS1 were specifically enriched in pathways related to immune activation, including platelet aggregation plug formation and were mostly up-regulated compared to immunosuppressive subtypes. Enriched HSF1 activation in CMS1 was also reported to orchestrate inflammation and ECM remodeling(48). *Flammeovirga, Sutterella*, and *Algiphilus* with increased abundance were positively correlated with genes from endothelin pathway, such as *COL1A2*, *COL3A1*, and *JUN*. *COL1A2* and *COL3A1* are collagen genes, which have been associated with cancer metastasis via regulating WNT pathway(49). *Desulfotalea*, a sulfate-reducing delta-Proteobacterium with putative cutC genes, was negatively associated with *LDLR* in plasma lipoprotein assembly remodeling and clearance pathway. In CMS2, the enriched host pathways were related to integrins, such as integrin1, integrin3 signaling. Our data highlighted that differentially depleted *Sutterella* was positively associated with a panel of genes in integrins-related signalings, such as *THBS2*, *PDGFRB*, and collagen genes *COL6A1* and *COL6A2*. Most of these genes were down-regulated compared to immune-high subtypes, which were consistent with the suppression of integrin3 pathway in CMS2(5). CMS3 was specifically enriched with biosynthetic and metabolic pathways, including glycosphingolipid biosynthesis lacto and neo lacto series, synthesis of bile acids and bile salts, sialic acid metabolism. Notably, *Methanomethylovorans* and *Metallosphaera*, two archaeal genera, were negatively associated with *FUT2*, *FUT3*, and *FUT6*. The knockdown of *FUT* genes can potentially inhibit the biosynthesis of certain oligosaccharide chains on tumor cell surface, making them desirable therapeutic targets(50). The abundance of *Ornithobacterium* was positively correlated with the gene expression of *SLC35A1*, a transporter of CMP-sialic acid, which modulates the immune system in diverse ways(51). In CMS4, specifically up-regulated host gene enriched pathways were mostly involved with ECM collagen construction, such as collagen biosynthesis and modifying enzymes, non-integrin membrane ECM interactions, assembly of collagen fibrils and other multimeric structures. Notably, *Sutterella*, *Collimonas*, and *Campylobacter* with increased abundance were positively associated with the majority of genes in collagen biosynthesis and modifying enzymespathway, such as *COL1A1*, *COL1A2*, *COL3A1*, *COL5A1*, *COL8A2*, and *CRTAP*. Moreover, the enhanced expression of *COL1A1* and *COL1A2* indicated the activation of wound healing CAFs, which was also a representative signature of CMS4(5, 8).

**Figure 5.**
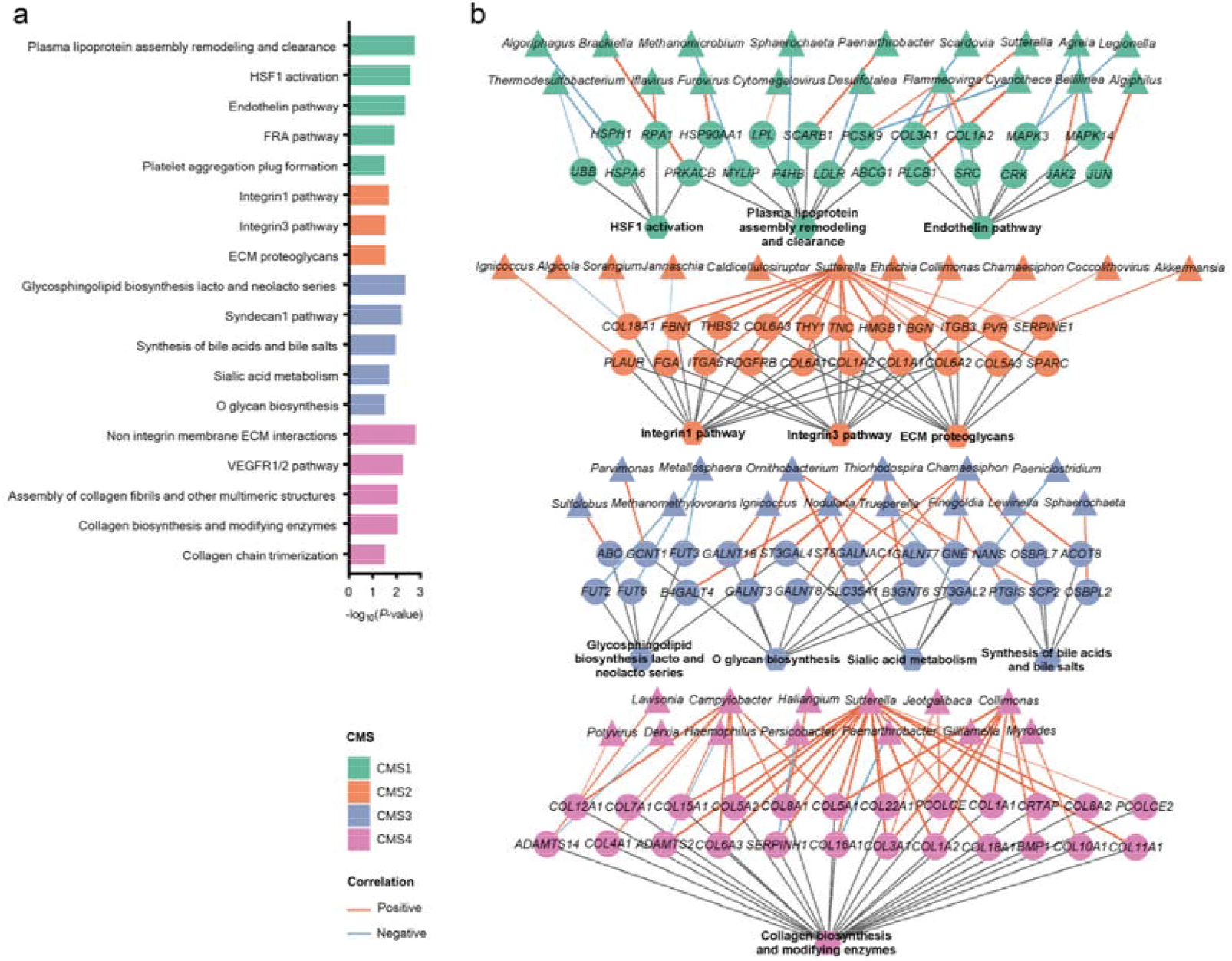
Interaction networks of host genes, gut microbiota, and enriched pathways of CMSs. **(a)**Boxplot of the pathways enriched with host genes that were significantly associated with microbes. Analyses were performed individually in each CMS(color coded). **(b)**The corresponding interaction network of (a), consisting of relevant host genes, gut microbiota, and enriched pathways. For nodes in the network, triangular nodes represent gut microbes, circular nodes represent host genes, hexagon nodes represent pathways. The colors of the nodes indicate different CMSs. The colors of the edges indicate positive (blue), negative (red), or gene-pathway(grey) associations while edge thickness represents Spearman rho coefficient.

Inspired by the distinct status of ferroptosis among CMSs, we then focused on ferroptosis-related gene-microbe associations, namely associations involving ferroptosis-related genes. We identified 35 ferroptosis-related gene-microbiota associations in CMS1, 7 associations in CMS2, 13 associations in CMS3, and 49 associations in CMS4(Figure 6). In CMS1, one-to-one associations were identified between ferroptosis-related genes and microbial genera. Among these, the most differentially enriched genus in CMS1, *Flammeovirga*, was positively correlated with ferroptosis driver gene *DDR2*(Figure 6a). In CMS2, *Apibacter* positively correlated with driver gene *PGRMC1* and negatively correlated with suppressor gene *SRC*(Figure 6b). On the contrary, in CMS3, a positive correlation between *Ruminococcus* with suppressor gene *RARRES2* and negative correlations between *Chamaesiphon* with driver genes *ACSL4* as well as *IDH1* were identified(Figure 6c). In CMS4(Figure 6d), the majority of the associations were between *Sutterella*, including driver genes *DDR2*, *WWTR1*, and *TIMP1*, suggesting the vital role of *Sutterella* in ferroptosis dysregulation in CMS4.

**Figure 6.**
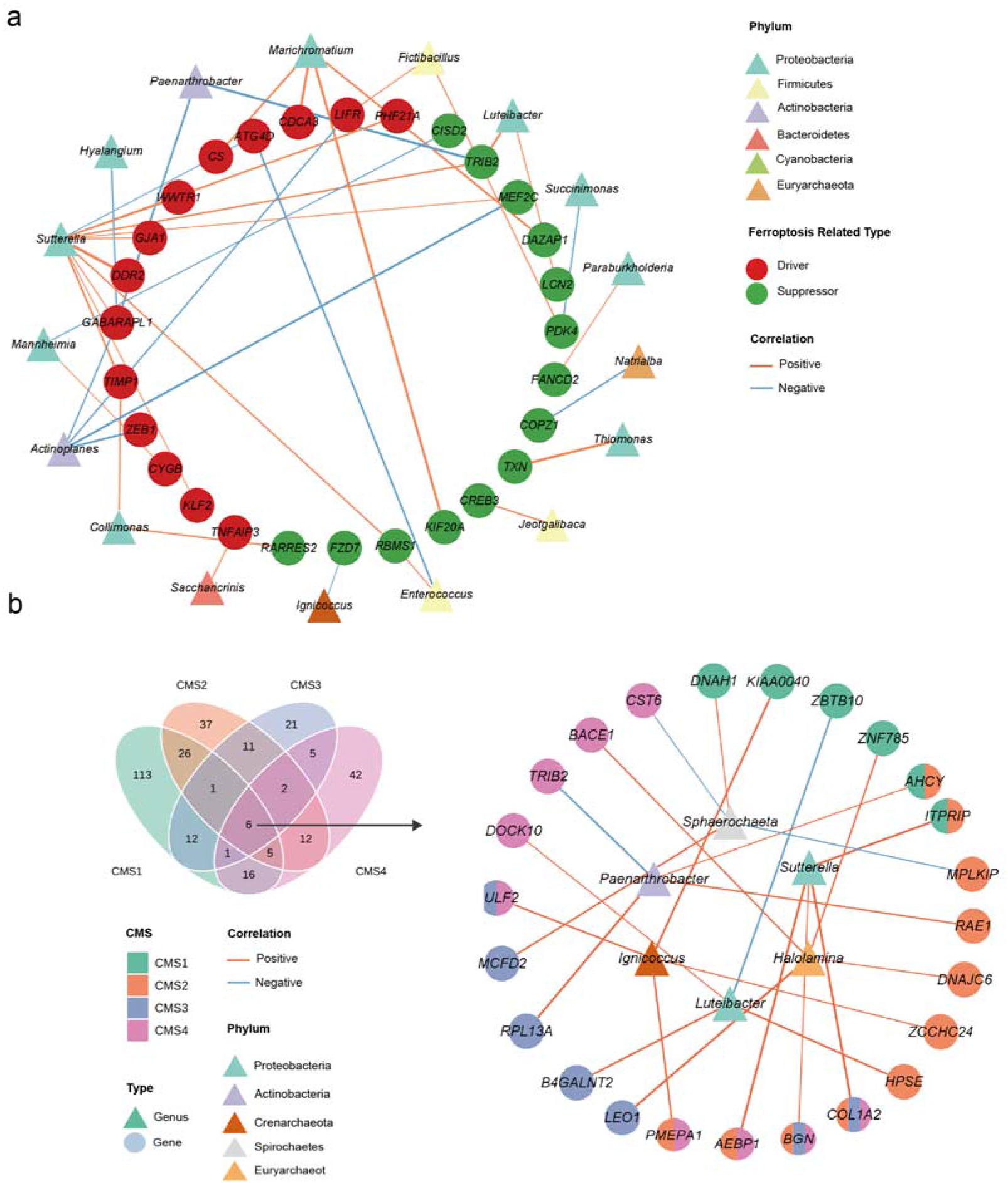
Ferroptosis-related gene-microbe associations in the CMSs. The gene-microbe associations involving ferroptosis-related genes in CMS1**(a)**, CMS2**(b)**, CMS3**(c)** and CMS4**(d)**. Triangular nodes represent gut microbes; circular nodes represent host genes. Colors of the triangles represent different phyla. The colors of the circles represent different functions: driver, suppressor or genes with multiple functions. The edge colors represent positive(blue) or negative(red) associations while edge thickness represents Spearman rho coefficient.

For a better understanding of the intratumoral microbes associated with host genes across CMSs, we examined the common genera mediating gene-microbiota associations in all CMSs. We identified six genera that appeared in all CMSs, that is *Sutterella*, *Paenarthrobacter*, *Luteibacter*, *Sphaerochaeta*, *Ignicoccus*, and *Halolamina*(Figure 7). These genera correlated with different genes. Among these correlations, *Sutterella*, *Luteibacter*, *Sphaerochaeta*, and *Halolamina* exhibited increased abundances in immune-high subtypes compared to immunosuppressive subtypes. *Sutterella* positively associated with different genes across CMSs, including *COL6A3* in CMS1, *GFPT2* in CMS2, *COL1A2* in CMS3 and *BGN*, *SERPING1* in CMS4. *Luteibacter* showed positive correlations with genes in immunosuppressive subtypes, such as heparinase(*HPSE*) in CMS2 and *B4GALNT2* in CMS3. *Paenarthrobacter* showed increased abundances in CMS2, with positive correlations with genes including *AHCY* in CMS1, *RAE1* in CMS2, and *RPL13A* in CMS3. Some negative associations were also identified. For example, *Luteibacter* negatively correlated with *ZBTB10* in CMS1 and *NEU1* in CMS4. *Paenarthrobacter* exhibited negative correlations with genes such as *TBRG4* and *WDR7* in CMS4.

**Figure 7.**
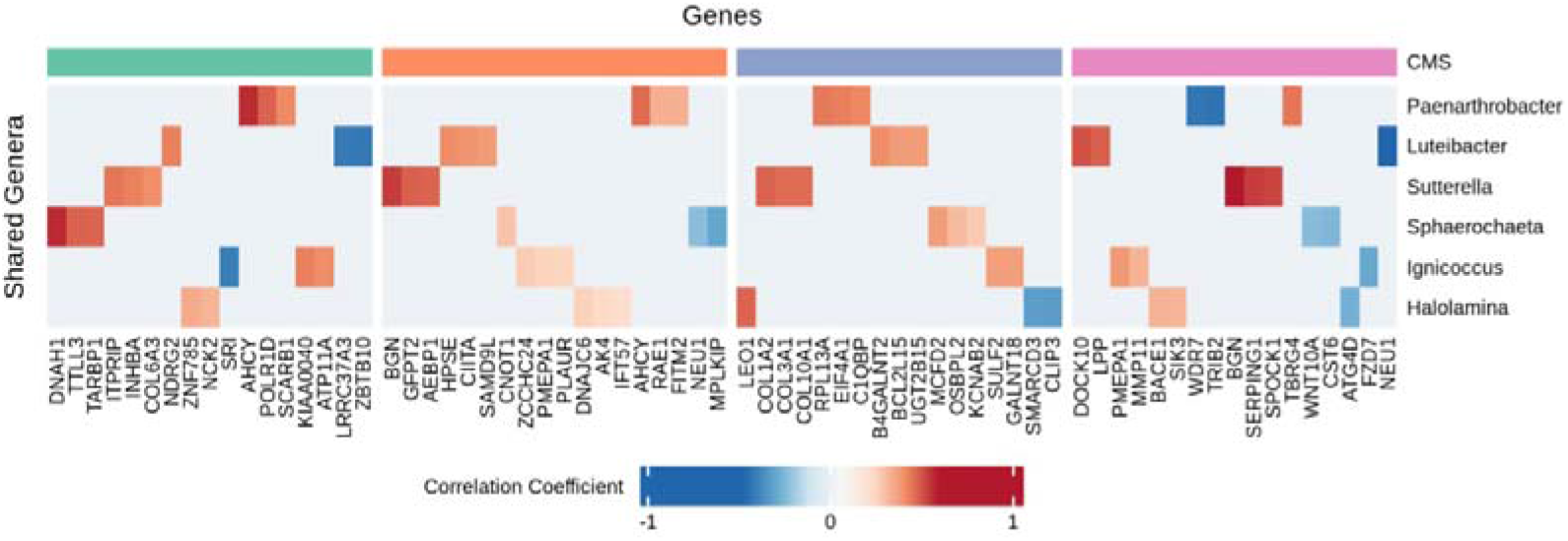
Heatmap of the gene-microbe associations of the genera common for all CMSs. Top associated genes of six common are plotted. Genes were clustered within each CMS.

## Discussion

In this proof-of-concept study, we systematically profiled the landscapes of the TME of each CMSs in CRC. We depicted the distinctions of host genes and immune environments, especially the ferroptosis-related genes. Host transcriptomes, intratumoral microbial compositions, and ecological communities across all CMSs were characterized. Notably, CMS-specific host gene-microbiota association patterns were profiled for the first time.

Microbiota dysregulation in CRC hasbeen broadly acknowledged based on fecal 16S rRNA and whole metagenome sequencing. Microbial biomarkers, including bacterial species and multi-kingdom species, are now emerging as potential non-invasive diagnostic and prognostic tools(15, 16, 19).Recently, tissue microbiota hasbeen detectedin tumors and implicated in regulating TME, including inflammatory mediators, resident and recruited immune cells(52, 53). As for CRC, Younginger et al has highlight that the associations between the intratumoral microbiota and host gene among CMSs were both species- and tumor-context-specific, for example, collagen-related pathways were associated with *P. dorei* in CMS2 and *F.animalis*in CMS4, respectively(54). However, the explorations of whole landscape of intratumoral microbiota are still in infancy, and the pan-cancer intratumoral microbiome data generated by Poore et al from RNA-Seq studies(20) provided an essential resource for tumor researches. We found that the intratumoral microbiota in CRC was dominated by Proteobacteria accounting for 38.7-38.8%. Besides, archaea and viruses were also observed in CRC tissues(Figure 4b). Considerable differences were identified among CMSs in intratumoral microbial composition. SCFA-producing genera were increased in CMS1 but decreased in CMS2, such as *Akkermansia*, *Lachnoclostridium*, and *Ruminococcus*(See Supplementary Table 3a and 3b). Alterations in SCFA levels could impact colonic health and predispose colonocytes to aberrant metabolism and tumor transformation(55). Therefore, such contrast in SCFA-producers in different CMSs highlighted the heterogeneity of intratumoral microbiota and indicates distinct host-microbiota crosstalk in different CMSs. Besides, the immune-high subtypes exhibited an opposing abundance pattern in intratumoral microbiota against immunosuppressive subtypes, with top differentially enriched genera in immune-high subtypes such as *Flammeovirga*, *Sutterella* and *Collimonas* depleted in CMS2 and CMS3(See Supplementary Table 3b and 3c). Among these genera, *Flammeovirga* was positively associated with the infiltrated CD8^+^ T cells(56), in line with the higher fraction of CD8^+^ T cells in CMS1 in our study. *Sutterella* was found to be the most abundant genus in colon adenocarcinomas(57, 58). Although it induced mild or even negligible inflammatory responses, *Sutterella* could serve as an IgA-degrading bacteriathat results in homeostasis disruption and a TME conducive topathobiont invasion(59). Furthermore, the intratumoral microbial co-abundance associations in each CMS exhibited disparate patterns. The networks of CMS1 and CM4 were dominated by Firmicutes and Proteobacteria genera compared to the other CMSs. *Enterococcus* from Firmicutes, a hub genus in CMS4, could stimulate *CXCL10* and thus lead to inflammation and take part in CRC development(60). The strain of *Enterococcus*, *E. faecalis*, a double-edged sword in CRC, attenuates inflammation via T helper(Th)-1 and Th17 suppression while accelerates EMT via macrophage *MMP9* activation(61). Consistently, a higher expression of in *CXCL10* and *MMP9* was observed in immune-high subtypes(See Supplementary Table 2a and 2d). The infection of *Helicobacter* in phylum Proteobacteria, a hub genus ranked sixth in CMS1, could induce pro-inflammatory responses in the intestine of mouse models, accompanied by a reduction of Treg cells(62). Similarly, our study revealed that the amount of Treg cells was decreased in CMS1(Figure 3b).

We identified gene-microbe associations through LASSO panelized regression model and performed pathway enrichment based on genes in these associations in each CMS. The interactions between host genes, tissue microbiota, and biological pathway collectively revealed discordant biological mechanisms in each CMS. At the pathway level, CMS1 interaction network was featured with inflammation related pathways while CMS2 exhibited suppression of integrins pathways. CMS3 and CMS4 interaction networks were populated by biosynthesis and metabolism pathways and ECM-related pathways, respectively. Likewise, we also identified CMS-specific gene-microbe associations. Although some genera appeared in multiple CMS interaction networks, they were associated with different host genes in each CMS. For example, the abundance of *Sutterella* was correlated with host genes in integrin signaling in CMS2 but with collagen-related genes in CMS4(8). Similarly, the abundance of *Sphaerochaeta*, which was reported to have notable enrichment in CRC and inflammatory bowel diseases(IBD)(63, 64), correlated with *P4HB* in CMS1 but with *OSBPL2* in CMS3.

Recently, Luo et al provided a comprehensive TME landscape mediated by ferroptosis in CRC, revealing that ferroptosis was positively correlated with CMS1 and CMS4 characteristics and might affect CRC through immune activation and stromal pathways activation(44). Consistently, our study also found that immune-high subtypes showed ferroptosis activation compared to the other CMSs(See Supplementary Figure 2-5). Moreover, we examined the ferroptosis relevant gene-microbe associations and discovered more complex association networks in immune-high subtypes(Figure 6), which suggests that tissue microbes might be significant contributors to ferroptosis dysregulation in CMS1 and CMS4. For example, ferroptosis related genes *CISD2* and *GCH1* are involved in immune activation and CD8^+^ T inflitration(44). Correspondingly, we observed the highest level of CD8^+^ T cells and associations between *CISD2* and *Sphaerochaeta*, and between *GCH1* and *Alphapapillomaviru* in CMS1 (Figure 6a). Other ferroptosis related genes such as *PLIN4* and *HIC1* are involved in stromal infiltration(44), and in CMS4, *PLIN4* was associated with *Bibersteinia,* while *HIC1* was associated with *Sutterella*, *Haliangium*, and *Mannheimia*(Figure 6d). Besides, *Sutterella* served as the key genus in ferroptosis-related gene-microbes network in CMS4(Figure 6d), suggesting that it might affect CRC through ferroptosis dysregulation.

In conclusion, we systematically described the CMS-specific landscapes of TMEs encompassing multiple aspects, including the expression of host genes, immune infiltrations, tissue microbiota, and biological pathways based on TCGA-CRC RNA-Seq studies. Especially, we first profiled the CMS-specific gene-microbe associations, revealing distinct interaction patterns in each CMS and further exploredthe potential functional relevance in CRC pathophysiology. The growing understanding of heterogeneity in gene-microbe associations across CMSs could shed light on novel mechanisms in CRC development and potential therapeutic targets.

## Supplementary Figure Titles and Legends

**Supplementary Figure 1.** Principal component analysis(PCA) across sequencing platforms. **(a)** PCA of host gene expression data before ComBat correction. **(b)** PCA of host gene expression after ComBat correction. **(c)** PCA of gut microbial abundance data before Voom-SNM correction. **(d)** PCA of gut microbial abundance data after Voom-SNM correction.

**Supplementary Figure 2.** Heatmap displaying the co-expression of ferroptosis-related CMS1-specific genes. Genes in row(column) were classified as driver(red), suppressor(green), unclassified(grey). Genes that were assigned to multiple groups were colored in blue.

**Supplementary Figure 3.** Heatmap displaying the co-expression of ferroptosis-related CMS2-specific genes. Genes in row(column) were classified as driver(red), suppressor(green), unclassified(grey). Genes that were assigned to multiple groups were colored in blue.

**Supplementary Figure 4.** Heatmap displaying the co-expression of ferroptosis-related CMS3-specific genes. Genes in row(column) were classified as driver(red), suppressor(green), unclassified(grey). Genes that were assigned to multiple groups were colored in blue.

**Supplementary Figure 5.** Heatmap displaying the co-expression of ferroptosis-related CMS4-specific genes. Genes in row(column) were classified as driver(red), suppressor(green), unclassified(grey). Genes that were assigned to multiple groups were colored in blue.

**Supplementary Figure 6.** Differences in the infiltration of the remaining 14 immune cells calculated by CIBERSORTx among CMSs.

**Supplementary Figure 7.** The microbial diversity among CMSs. (a)Comparison of alpha diversity (Gini Simpson index) among CMSs. (b) Principal coordinate analysis (PCoA) of all samples based on Bray–Curtis distance, estimating the beta diversity of CMSs.

**Supplementary Figure 8.** Procrustes analysis and mantel test between host gene expression and gut microbial abundance data. Aitchison’s distance was used for host gene expression data and Bray-Curtis distance was used for gut microbial abundance data (green, triangles).

## Supplementary Table with Titles and Legends

**Supplementary Table 1**: The Distribution of CMS Cohorts by Clinical Characteristics.

**Supplementary Table 2**: Differential and KEGG enrichment analysis of host gene expression. Related to Figure 2. **(a)**Differential Genes of CMS1. **(b)**Differential Genes of CMS2.**(c)**Differential Genes of CMS3. **(d)**Differential Genes of CMS4. **(e)**The Enriched KEGG Pathways of Differential Genes of CMS1(P value<0.05). **(f)**The Enriched KEGG Pathways of Differential Genes of CMS2(P value<0.05). **(g)**The Enriched KEGG Pathways of Differential Genes of CMS3(P value<0.05). **(h)**The Enriched KEGG Pathways of Differential Genes of CMS4(P value<0.05).

**Supplementary Table 3**: Differential and co-abundance analysis of gut microbiome. Related to Figure 4.**(a)**Differential Genera of CMS1. **(b)**Differential Genera of CMS2. **(c)** Differential Genera of CMS3. **(d)** Differential Genera of CMS4. **(e)**CMS1-specific Co-abundance Network Nodes. **(f)**CMS2-specific Co-abundance Network Nodes. **(g)**CMS3-specific Co-abundance Network Nodes. **(h)**CMS4-specific Co-abundance Network Nodes.

**Supplementary Table 4**: Gene-microbe associations and the enriched pathways of host genes from gene-microbe associations. **(a)**CMS1-specific Gene-Microbe Associations. **(b)**CMS2-specific Gene-Microbe Associations. **(c)**CMS3-specific Gene-Microbe Associations. **(d)**CMS4-specific Gene-Microbe Associations. **(e)**The Enriched Pathways of Host Genes from CMS1-specific Gene-Microbe Associations. **(f)**The Enriched Pathways of Host Genes from CMS2-specific Gene-Microbe Associations. **(g)**The Enriched Pathways of Host Genes from CMS3-specific Gene-Microbe Associations. **(h)**The Enriched Pathways of Host Genes from CMS4-specific Gene-Microbe Associations.

