## Supplementary material for "Host-microbiota interactions contributing to the heterogeneous tumor microenvironment in colorectal cancer": Supplemetary Figure1-8

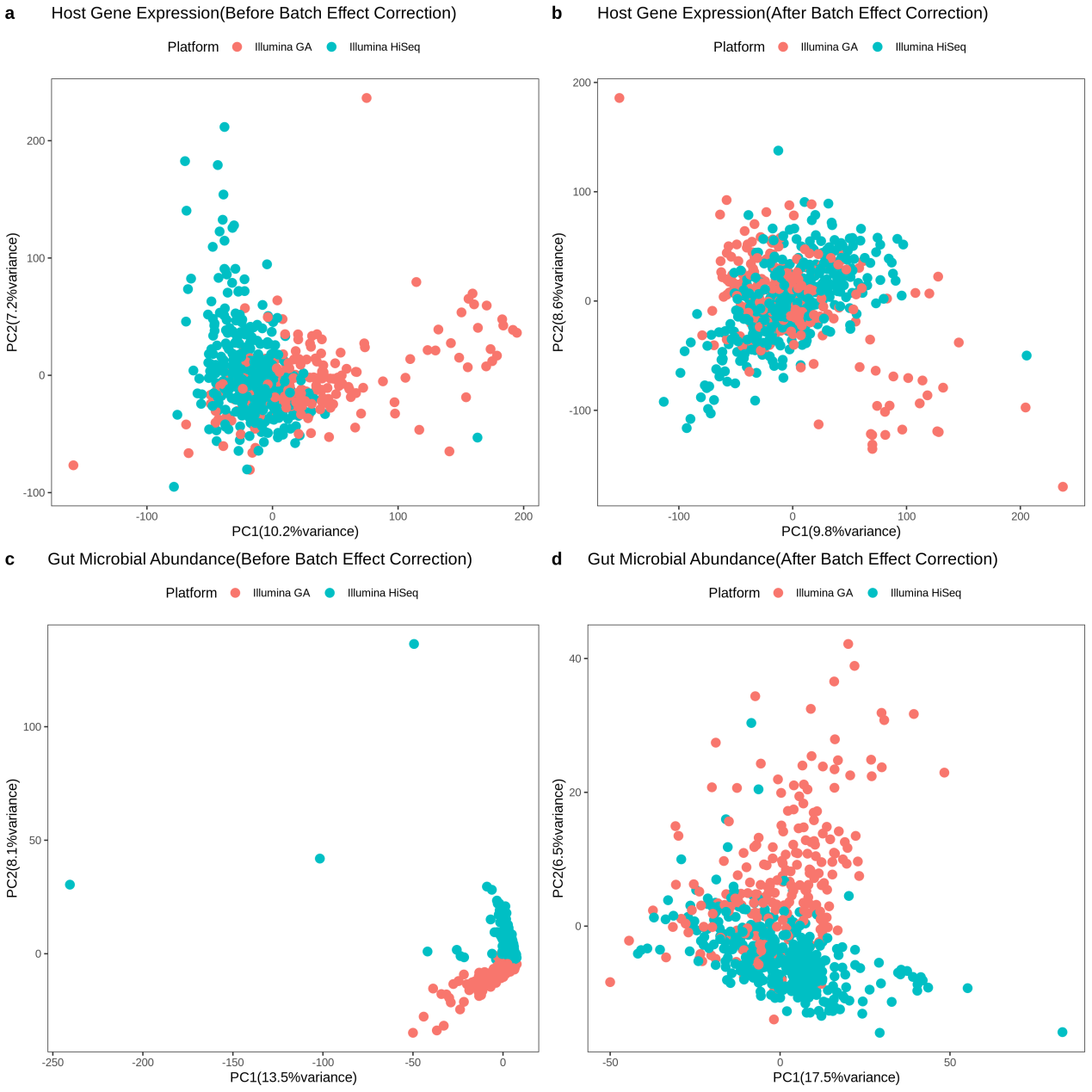

**Supplementary Figure 1**. Principal component analysis(PCA) across sequencing platforms. **(a)** PCA of host gene expression data before ComBat correction. **(b)** PCA of host gene expression after ComBat correction. **(c)** PCA of gut microbial abundance data before Voom-SNM correction. **(d)** PCA of gut microbial abundance data after Voom-SNM correction.

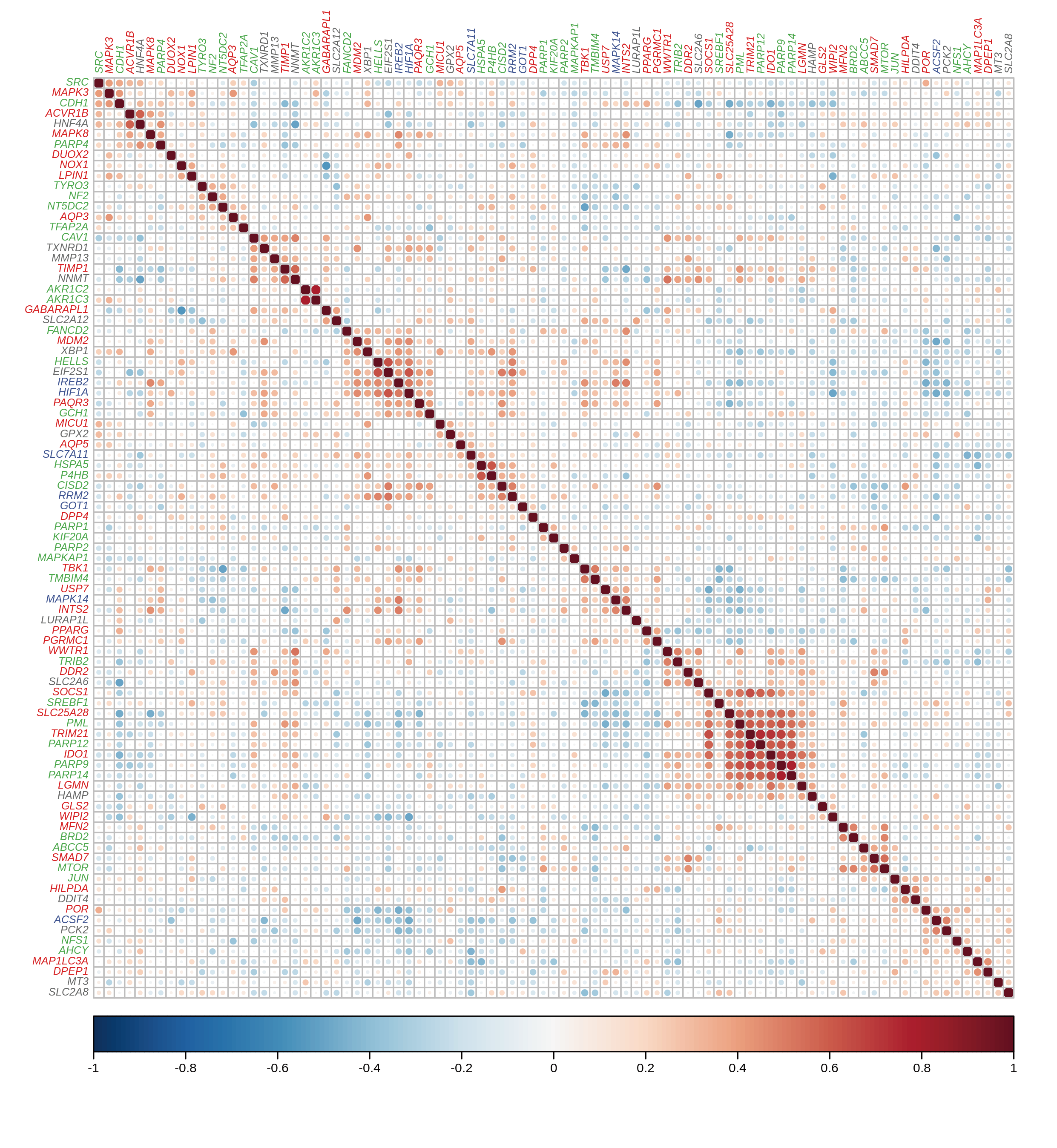

**Supplementary Figure 2**. Heatmap displaying the co-expression of ferroptosis-related CMS1-specific genes. Genes in row(column) were classified as driver(red), suppressor(green), unclassified(grey). Genes that were assigned to multiple groups were colored in blue.

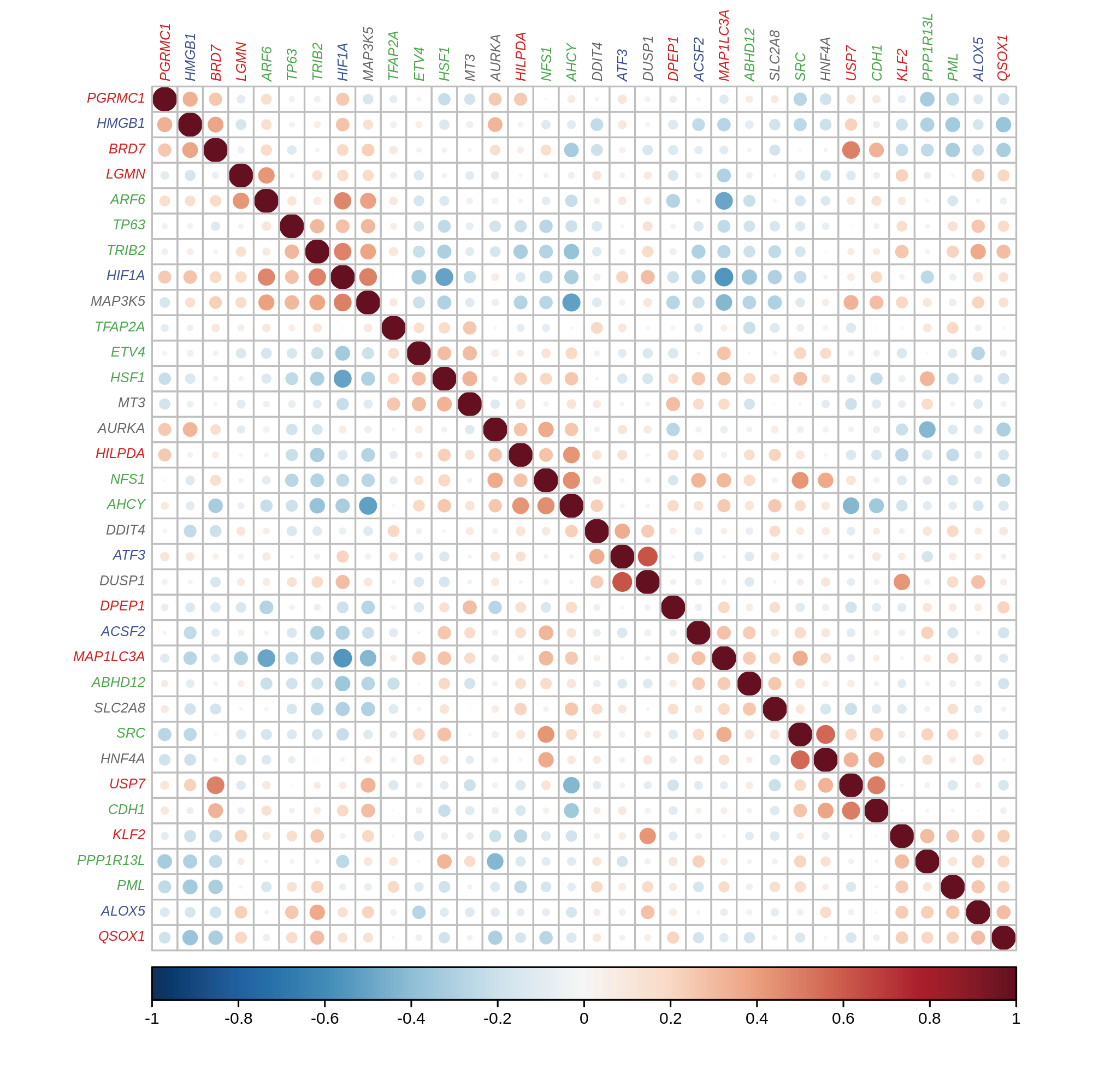

**Supplementary Figure 3**. Heatmap displaying the co-expression of ferroptosis-related CMS2-specific genes. Genes in row(column) were classified as driver(red), suppressor(green), unclassified(grey). Genes that were assigned to multiple groups were colored in blue.

**Supplementary Figure 4**. Heatmap displaying the co-expression of ferroptosis-related CMS3-specific genes. Genes in row(column) were classified as driver(red), suppressor(green), unclassified(grey). Genes that were assigned to multiple groups were colored in blue.

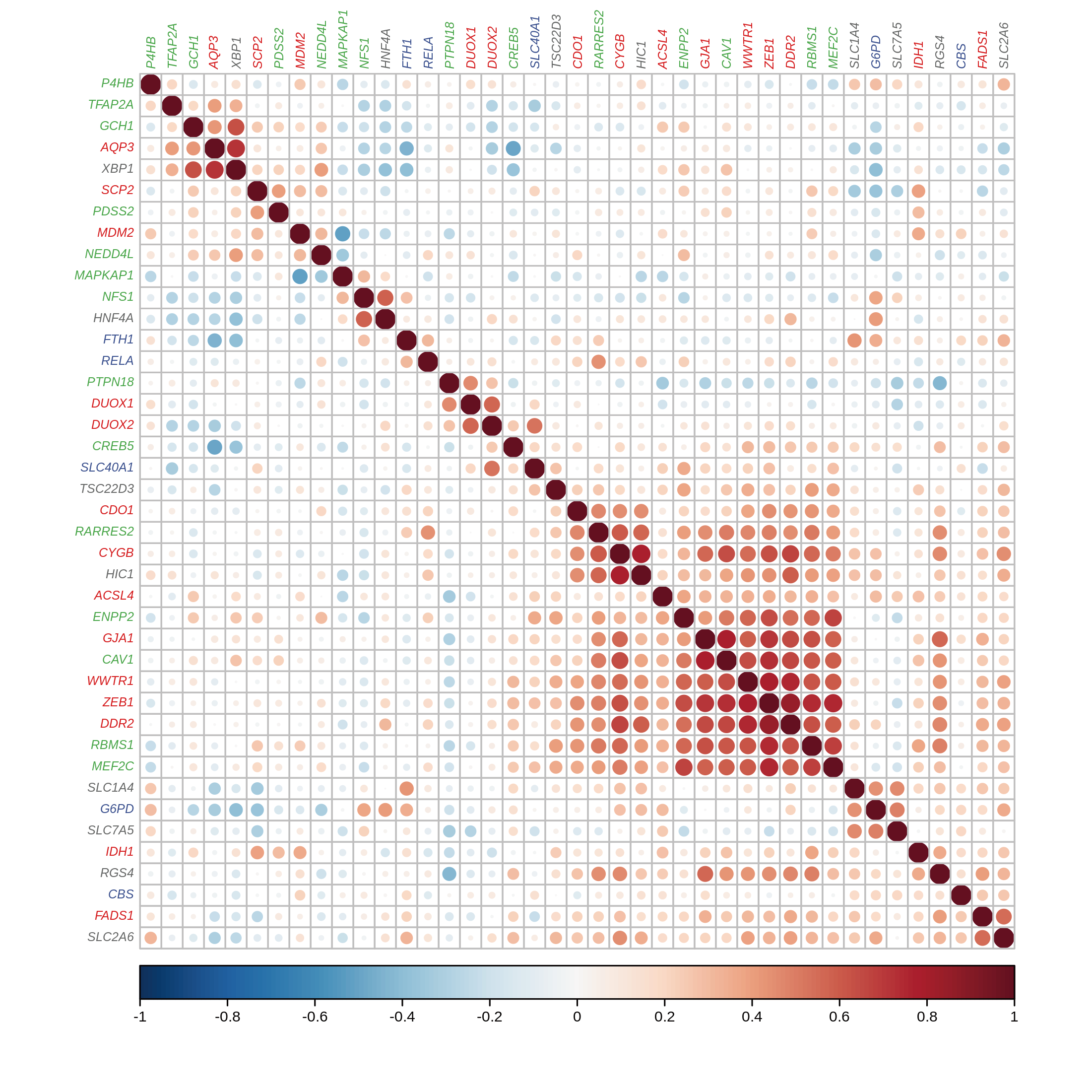

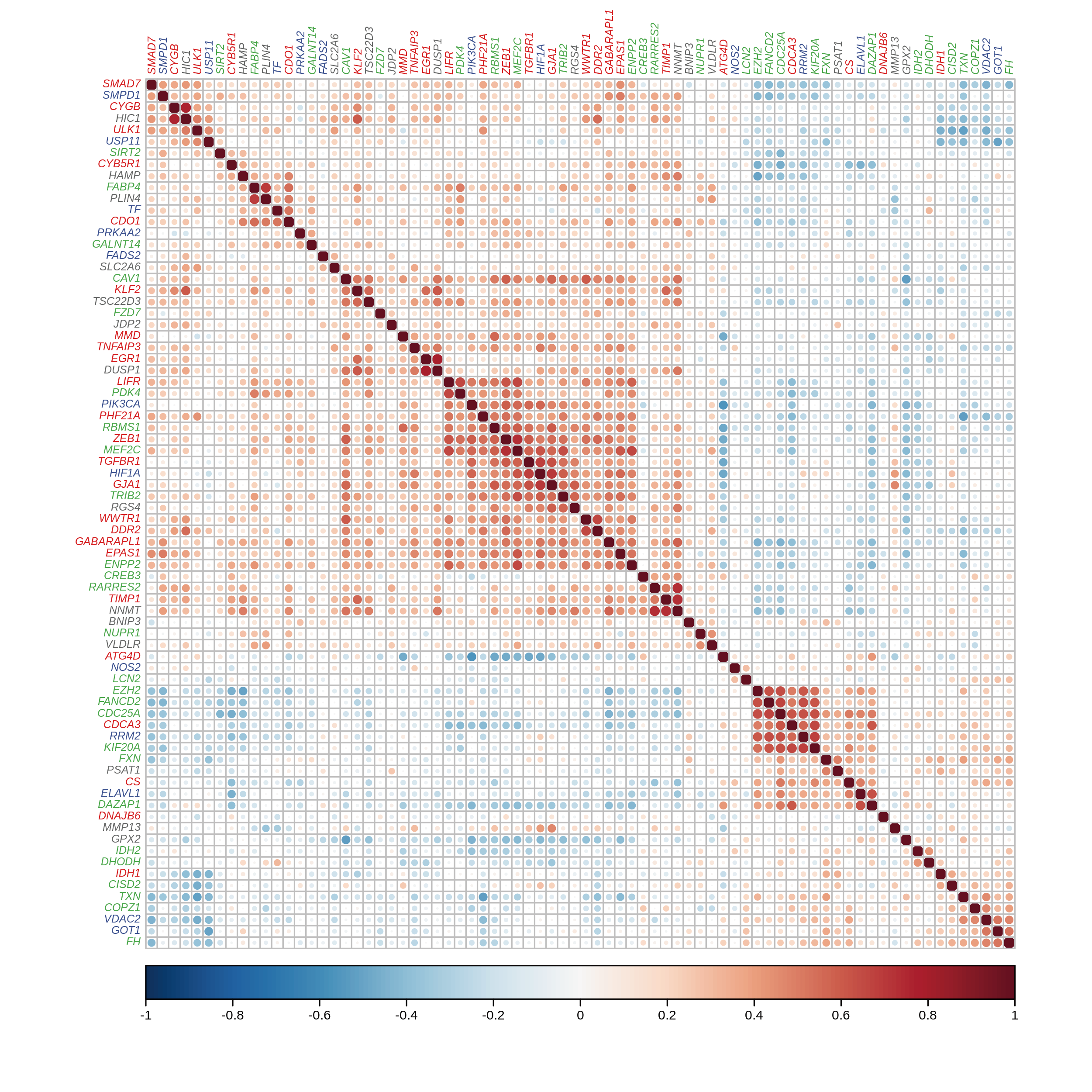

**Supplementary Figure 5**. Heatmap displaying the co-expression of ferroptosis-related CMS2-specific genes. Genes in row(column) were classified as driver(red), suppressor(green), unclassified(grey). Genes that were assigned to multiple groups were colored in blue.

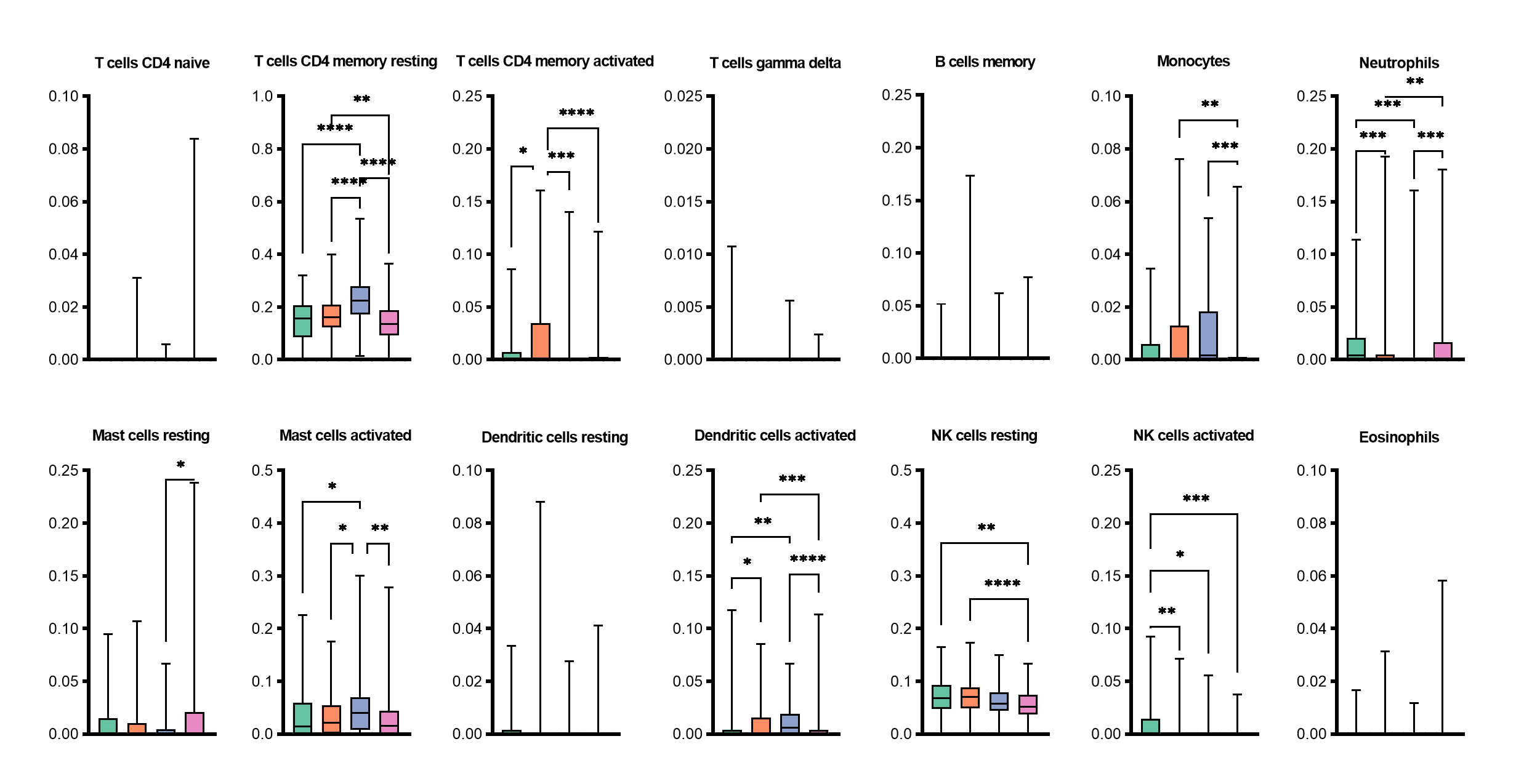

**Supplementary Figure 6**. Differences in the infiltration of the remaining 14 immune cells calculated by CIBERSORTx among CMSs.

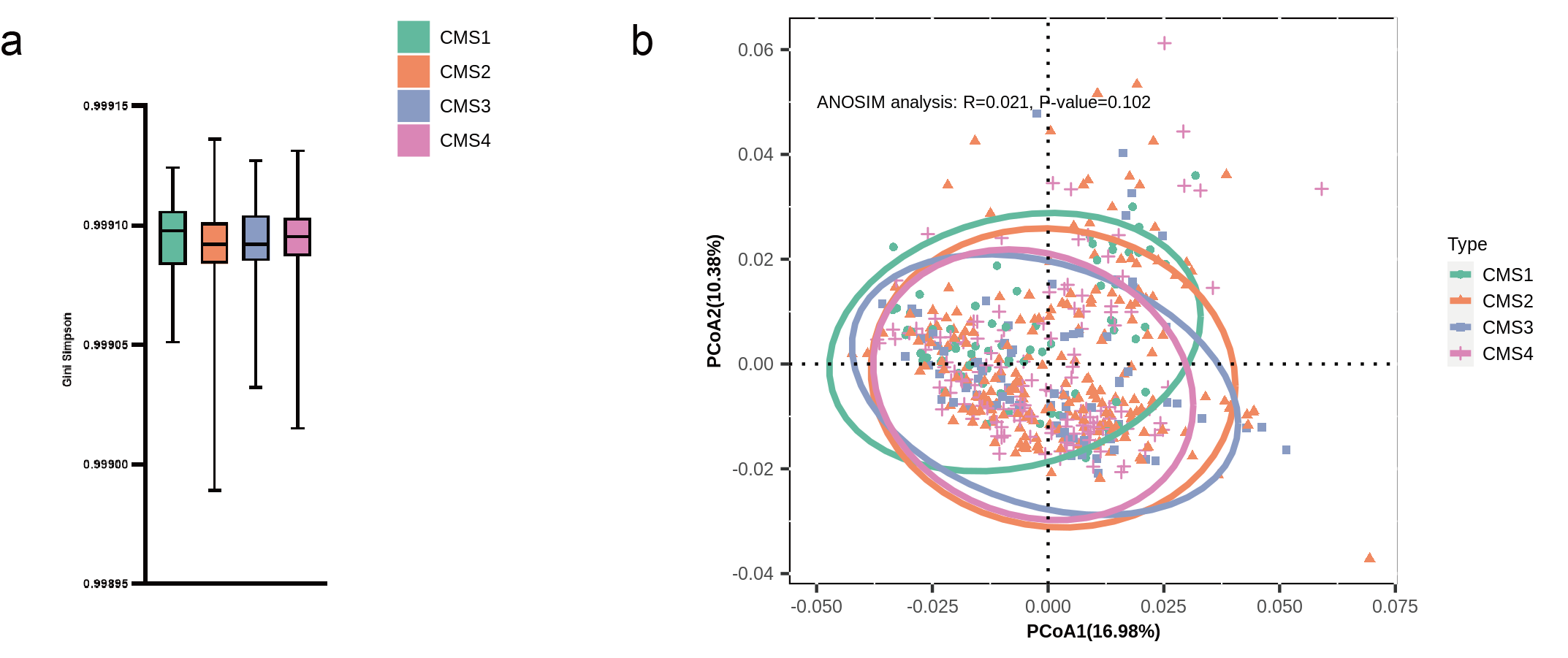

**Supplementary Figure 7**. The microbial diversity among CMSs. **(a)**Comparison of alpha diversity (Gini Simpson index) among CMSs. **(b)** Principal coordinate analysis (PCoA) of all samples based on Bray–Curtis distance, estimating the beta diversity of CMSs.

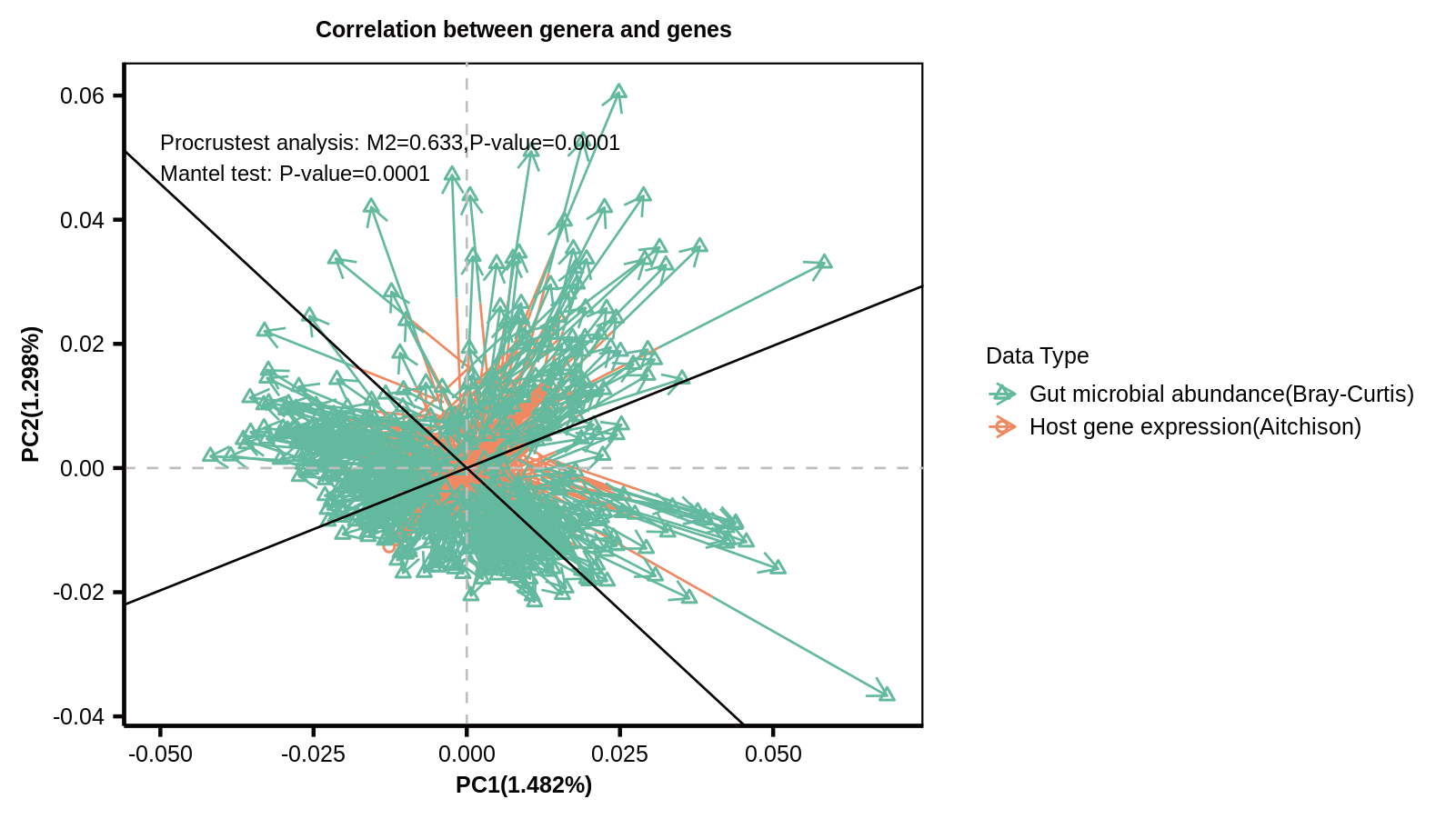

**Supplementary Figure 8**. Procrustes analysis and mantel test between host gene expression and gut microbial abundance data. Aitchison’s distance was used for host gene expression data and Bray-Curtis distance was used for gut microbial abundance data (green, triangles).
